# Large scale laboratory evolution uncovers clinically relevant collateral antibiotic sensitivity

**DOI:** 10.1101/2025.02.07.637158

**Authors:** Farhan R. Chowdhury, Veronica Banari, Vlada Lesnic, George G. Zhanel, Brandon L. Findlay

## Abstract

The increasing prevalence of antibiotic resistance is a critical challenge, necessitating the development of strategies to mitigate the evolution of resistance. Collateral sensitivity (CS)-based sequential therapies have been proposed to mitigate resistance evolution. However, the evolutionary repeatability of CS across different experimental conditions and its clinical relevance remain underexplored, hindering its potential for translation into clinical practice. Here, we evolve 20-24 lineages of *E. coli* against tigecycline (TIG) and piperacillin (PIP), antibiotics suggested to produce CS, through three separate laboratory adaptive evolution (ALE) platforms to test for the robustness of CS interactions and the effect of the choice of ALE on CS evolution. We generate over 130 resistant mutants and 540 resistance and collateral sensitivity measurements to identify a CS relationship between TIG and polymyxin B (POL) that is highly repeatable across all the ALEs tested, suggesting that this CS interaction is preserved across different evolution microenvironments. We determine the mechanism of this novel CS by showing that cells resistant to TIG deactivate the Lon protease and overproduce negatively charged exopolysaccharides, which in turn attracts the polycationic POL and renders cells hypersensitive to the drug. We find that this CS relationship is present in a clinical dataset of over 750 uropathogenic MDR *E. coli* isolates, and show that the soft agar gradient evolution (SAGE) platform best predicts collateral effects (CS, neutrality or cross resistance) in this dataset. Our study provides a framework for identifying robust CS with clinical implications that can reduce the emergence of resistance to our existing antibiotics.

## Introduction

Antibiotic resistance is spreading at an alarming rate, claiming the lives of over 1.2 million people every year (*1*). Developing strategies to slow down resistance evolution has become essential to combat antibiotic resistance (*2*). Sequential antibiotic therapy, where antibiotics are administered in a chronological sequence in an individual patient, has emerged as a potential strategy to slow down the evolution of resistance (*3*). In principle, this strategy interrupts the selection of resistant populations by changing the selective pressure via switching treatment to a different drug (*4*). Evolutionary trade-offs like collateral sensitivity (CS) have been proposed to improve the success of such sequential therapies by limiting the rate of bacterial evolution (*5–7*).

We now have numerous studies that have described large networks of collateral sensitivities in different bacteria (*6, 8–11*). However, investigations on their evolutionary repeatability via large scale experimental evolution are scarce, but are essential for successful clinical application (*12*). In addition, different laboratories use different adaptive laboratory evolution (ALE) platforms that are difficult to standardize (*6, 8, 9, 13, 14*). Since evolutionary outcomes can vary depending on the bacterial microenvironment (*12, 15*), it is important to determine the effect of the choice of the ALE platform on CS evolution.

In this study, we evolve 20-24 lineages of *Escherichia coli* to screen for CS between four drug pairs reported to exhibit CS, using three different ALE platforms widely used to study evolution and CS (*16*). We find that serial transfer and gradient plating-based ALE platforms agree well on the frequencies of CS, cross-resistance (CR) and resistance levels. However, the soft agar gradient evolution (SAGE) platform produces substantially lower frequencies of CS and higher incidence of CR compared to the other two platforms. To test the relevance of these CS/CR predictions from the different ALE platforms, we analyze antimicrobial susceptibility data from over 750 clinical uropathogenic multidrug resistant (MDR) *E. coli* strains to test for the presence of CS/CR relationships. We find that CS is almost entirely absent, but neutrality or CR is prevalent. However, we observe a significant association between increasing omadacycline (a third generation tetracycline) resistance and reduced colistin (polymyxin E) resistance.

Interestingly, out of the four drug pairs screened in our ALE experiments, SAGE showed significant CS in only one of them: a tigecycline (TIG) (a third generation tetracycline) and polymyxin B (POL) pair. Using genomics and phenotypic analysis, we describe, for the first time, the mechanism of polymyxin B CS in tigecycline-resistant bacteria. Our results highlight the power of large-scale ALE experiments in predicting repeatable CS relationships that hold potential to reduce resistance in MDR bacteria in the clinic.

## Results

### SAGE produces lower collateral sensitivities and higher cross resistances compared to other ALE platforms

We evolved 20-24 mutants against tigecycline (TIG) and piperacillin (PIP) separately using three different ALE platforms (*16*): SAGE (*17*), serial transfer liquid culture-based method (LQ) (*18*), and gradient plates (GP) (*19*) (Figure 1A). We also investigated CS profiles for nitrofurantoin (NIT) and ciprofloxacin (CIP), but were unable to achieve resistance at high enough frequencies in LQ (NIT, CIP) or GP (CIP) to generate the necessary sample sizes (Supplementary Table 1). SAGE also produced 8-16-fold increases in relative MIC (MIC of evolved lineage/MIC of WT) of TIG and PIP, while levels from LQ were limited to 2-4-fold increases (Figure 1B, C) before incurring frequent extinctions.

**Figure 1:**
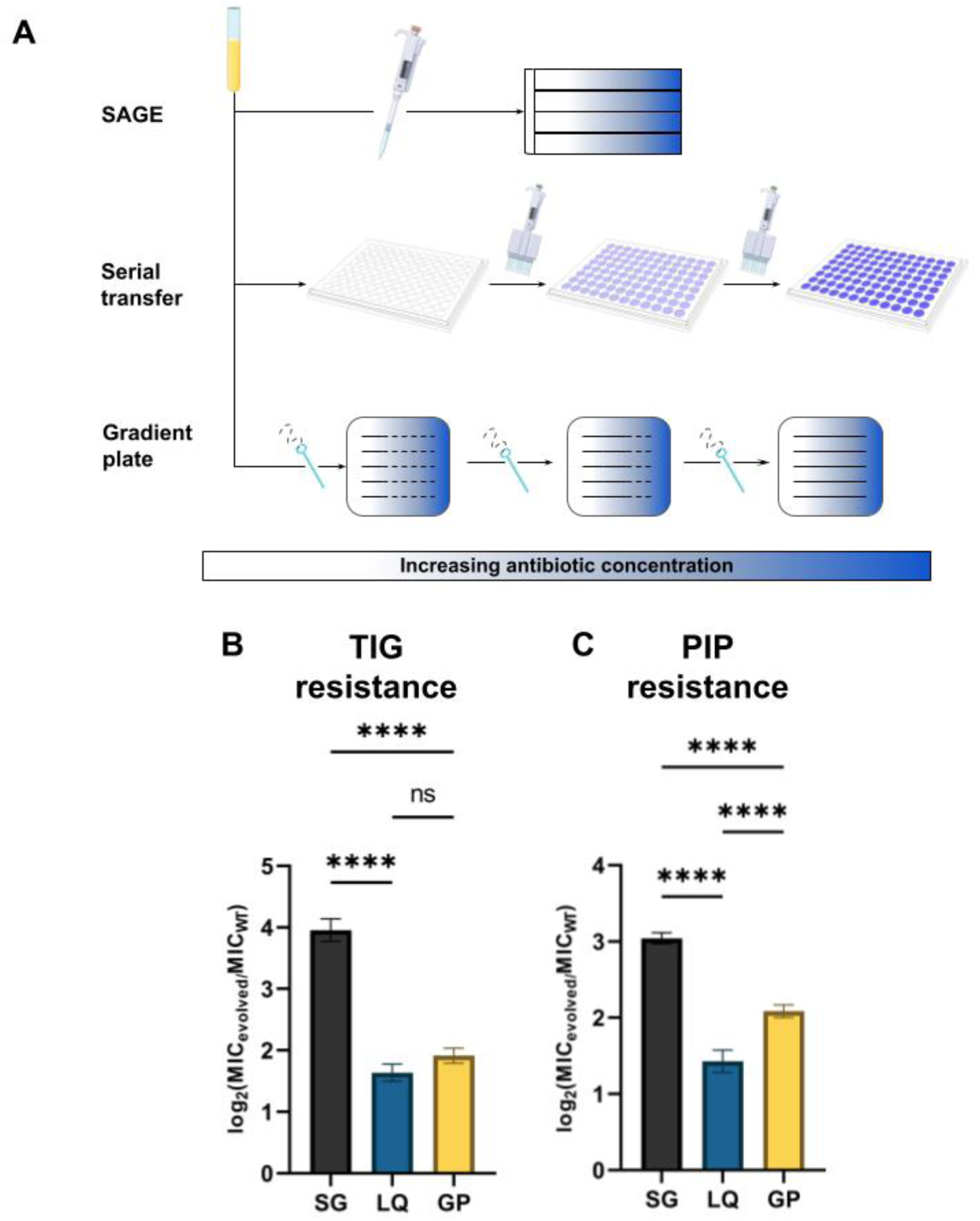
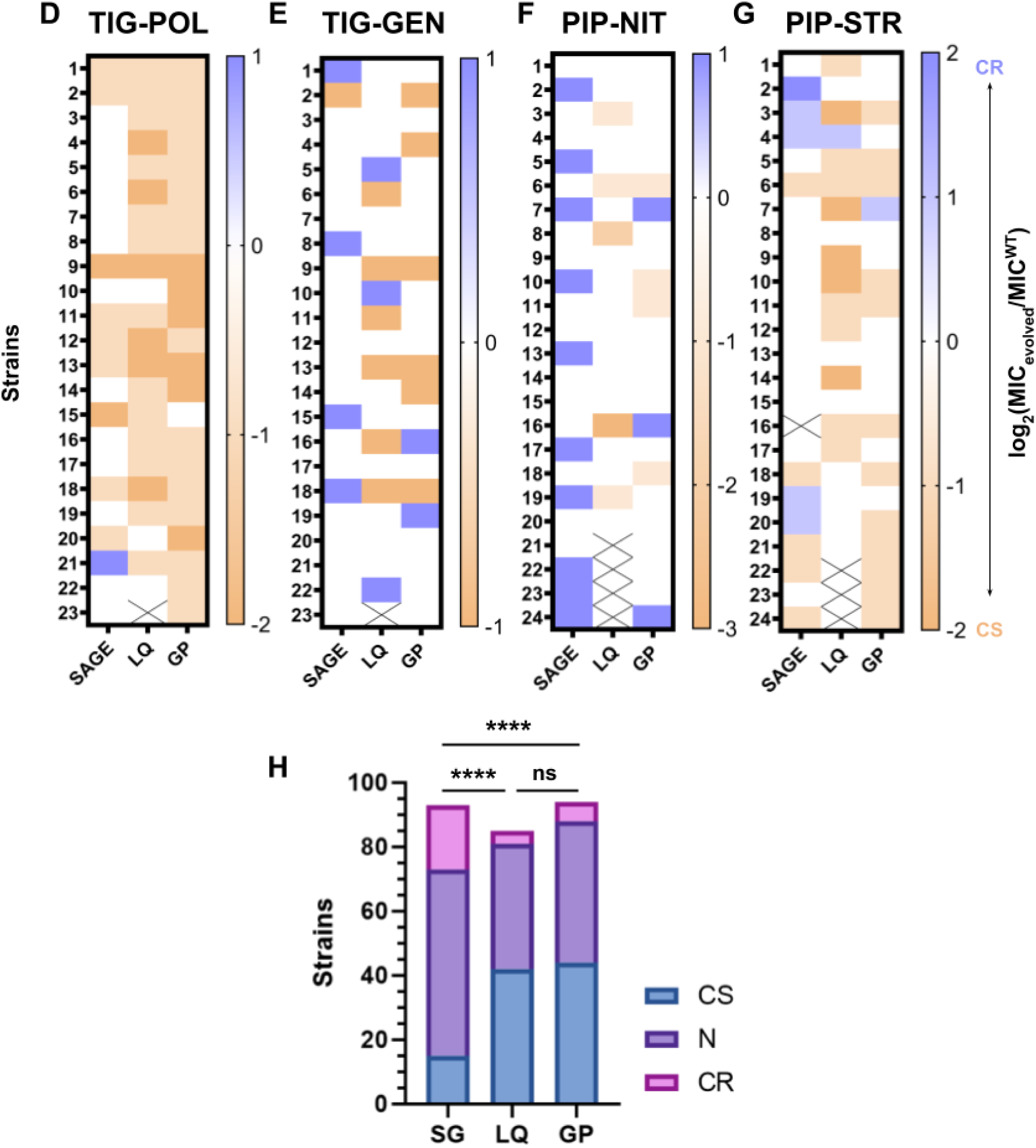
Evolution of antibiotic resistance using three different ALEs. (A) Schematic of the ALE platforms used to evolve resistance in this study. SG = SAGE. (B) Relative TIG and (C) PIP MICs of evolved mutants. ****p<0.0001, one-way ANOVA with Bonferroni correction. (D) - (G) CS profiles of mutants evolved to TIG and PIP. The labels on top show the antibiotics against which resistance was evolved and CS were measured. For example, “TIG-POL” denotes the POL CS measurements of TIG resistant mutants. (H) Combined CS, N and CR distributions from the three ALE platforms. SG = SAGE. ****p<0.0001, one-way ANOVA with Bonferroni correction. Statistical analyses were performed by comparing relative MICs of mutants from each platform.

Next, we screened for CS towards POL and GEN in the TIG-resistant lineages (*8*) and NIT and streptomycin (STR) in the PIP-resistant lineages (*6, 8*). A large proportion of TIG-resistant lineages exhibited CS to POL: ∼86% from LQ, ∼96% from GP, and ∼39% from SAGE (Figure 1D) (Supplementary Figure 1). We previously reported the presence of reciprocal CS between the POL-TIG pair (*20*). CS towards GEN was low in all platforms, with LQ and GP showing CS in ∼26% of lineages and SAGE in only ∼4% (Figure 1E) (Supplementary Figure 1). Cross resistance (CR) towards POL was almost entirely absent, and CR to GEN was present at very low frequencies (Figure 1D, E) (Supplementary Figure 1).

CS towards NIT was rare in the PIP-resistant lineages, present in only ∼17-25% of the LQ and GP strains and absent in the SAGE strains (Figure 1F) (Supplementary Figure 1). By contrast CR was common in the SAGE lineages, with ∼42% less susceptible to NIT (Figure 1F) (Supplementary Figure 1). About 50% of the PIP-resistant lineages from LQ and GP, and 21% from SAGE showed CS towards STR (Figure 1G) (Supplementary Figure 1). ∼20% of the SAGE mutants showed CR towards STR, but again this CR was rare in the other two platforms.

Overall, ∼16% of SAGE mutants showed CS compared to 49% and 47% from LQ and GP respectively (Figure 1H). Incidence of CR in SAGE mutants was much higher at about 21% than the 4-6% in LQ and GP mutants (Figure 1H).

### Tigecycline resistance evolves via similar pathways across ALE platforms

To compare the genomic adaptations of lineages evolved through the three different platforms and to identify differences in mutational profiles between strains that evolved CS and strains that did not, we whole genome sequenced 29 lineages adapted to TIG: nine from LQ (six with POL CS and three neutral), seven from GP (six with POL CS and one neutral) and 13 from SAGE (six with POL CS, six neutral, and one with POL CR) (Figure 2A). The genome profile of the lineage that showed POL CR from SAGE was a complete subset of the mutations appearing in the CS lineages, and was excluded from further analysis suspecting a two-fold random variation in MIC (*21*). Lineages acquired ∼1.2 mutations per strain from LQ, ∼1.7 strain from GP, and ∼1.5 mutations per strain from SAGE (Figure 2A). Strains from all platforms showed mutations in one or more of the following genes involved in TIG resistance: *lon*, *acrR*, and *marR* (Figure 2A) (Supplementary Figure 2A). Deactivation of the Lon protease, often achieved via mutations in its promoter region, spares the MarA, RamA and SoxS activators from degradation, increasing expression of genes that mediate resistance like *acrAB* (*22, 23*). Mutations in AcrR and MarR more directly relieve repression of the *acrAB* and *marRAB* operons, increasing efflux activity to drive TIG resistance (*23*). The primary method by which TIG resistance evolved is therefore through increased expression of efflux pumps, regardless of the evolutionary system.

**Figure 2:**
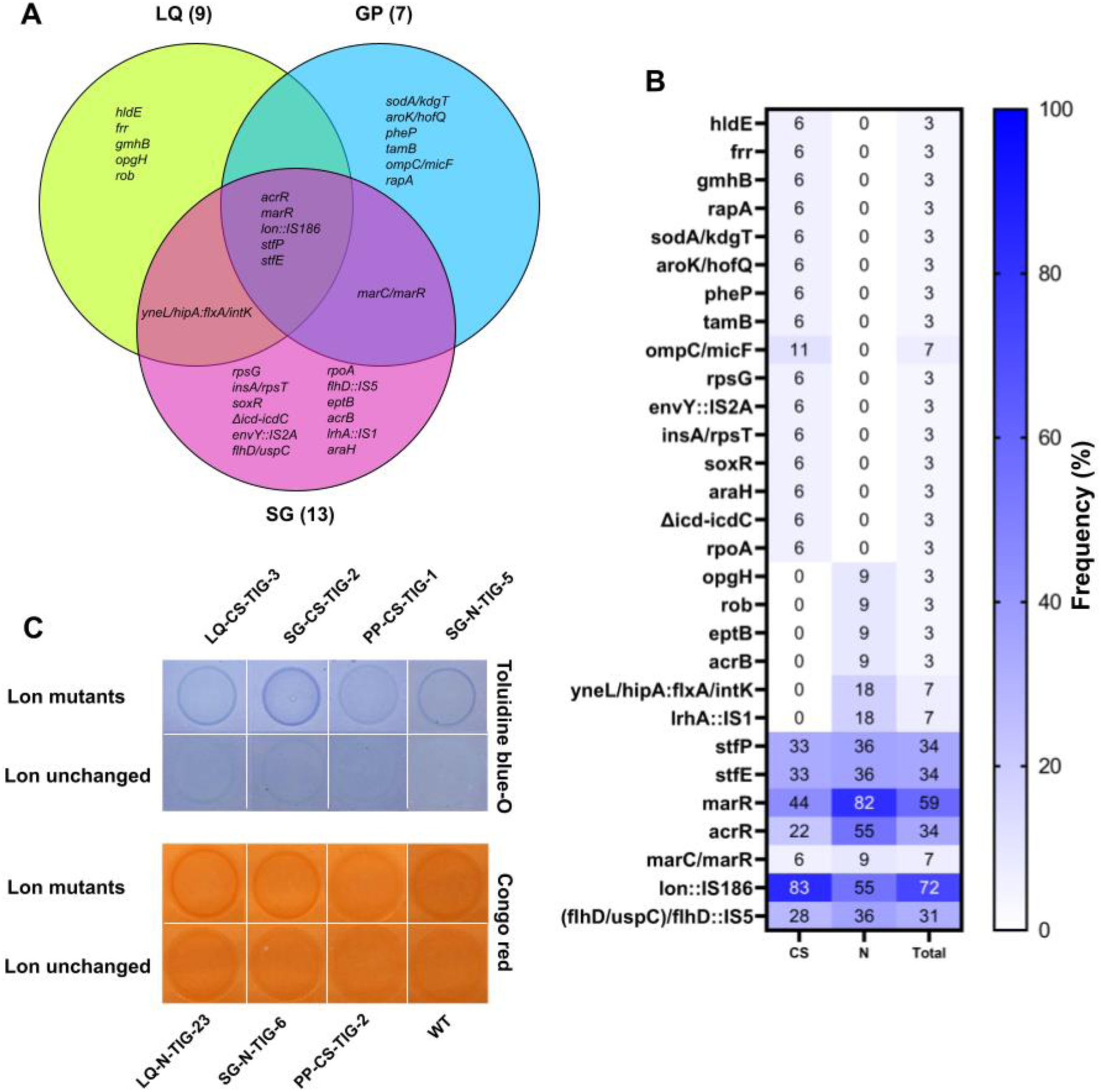
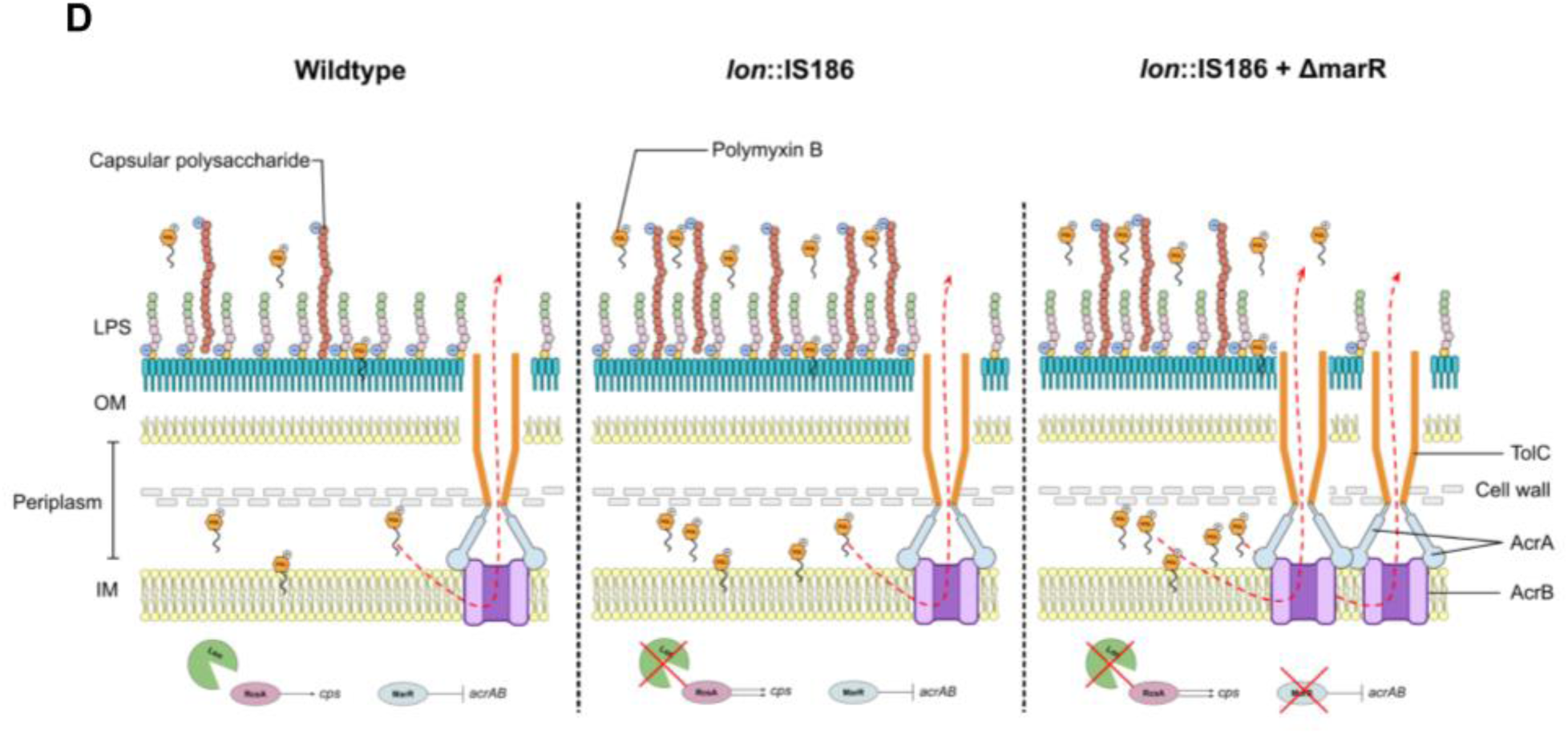
Genomic and phenotypic analysis reveals mechanism of POL CS in TIG resistant mutants. (A) Venn diagram showing common mutations between TIG mutants evolved from the three different platforms. The numbers in brackets denote the total number of mutants sequenced from that platform. (B) Frequency at which a mutation in the genes listed on the vertical axis appeared in all 29 mutants sequenced in this study, stratified by POL CS and N. “Total” denotes frequencies at which a mutation appeared among all strains. “sodA/kdgT” refers to an intergenic mutation between the genes *sodA* and *kdgT*. “envY::IS2A” denotes that gene *envY* was interrupted by the insertion element IS2A. “Δicd-icdC” refers to a deletion that affects all genes between and including *icd* and *icdC*. “yneL/hipA:flxA/intK” refers to a new junction between the intergenic regions of *yneL/hipA* and *flxA/intK*. Two different *flhD* mutations: (flhD/uspC) and flhD:IS5 were grouped together and reported as “(flhD/uspC)/flhD:IS5”. (C) Toluidine blue-O and congo red binding assays. The strain IDs of the Lon mutants are noted at the top of the toluidine blue-O panel, and IDs of the Lon unchanged mutants are noted at the bottom of the congo red panel. Strain IDs contain information about the platform used, phenotype and replicate number. For example, “LQ-CS-TIG-3” = the third TIG resistant replicate that showed CS to POL from the LQ platform. Strain SG-N-TIG-5 contained both the Lon and a MarR mutation. WT = wildtype. (D) The role of Lon and MarR mutations in POL CS. Left panel: WT cells. Lon degrades RcsA, limiting the expression of capsular polysaccharide genes (cps). MarR expression negatively regulates AcrAB expression. Middle panel: Lon deactivation spares RcsA from degradation, allowing expression of cps genes and causing the production of capsular polysaccharides. Increased negative charge on the membrane due to these polysaccharides causes increased accumulation of the polycationic POL, making cells more susceptible to POL. Right panel: MarR deactivating mutations allow increased AcrAB expression, increasing the number of efflux pumps that are able to pump out POL molecules, neutralizing POL CS.

### Polymyxin collateral sensitivity is linked to Lon protease deactivation

While POL CS in TIG-resistant *E. coli* has been previously reported (*8, 20*), the mechanism of POL CS remains unknown. Since CS was abundant in strains evolved through all three platforms, we hypothesized that mutation(s) that 1) occurred in all three platforms, 2) appeared frequently and 3) appeared more frequently in strains exhibiting CS may be responsible for POL CS. We tallied the frequencies of each mutation stratified by CS and neutrality (Figure 2B) (Supplementary Figure 2) and found that the *lon*::IS186 mutation, previously reported to be a mutation hotspot (*24*), was the only mutation that met these criteria (Figure 2B) (Supplementary Figure 2).

Cells that lack Lon activity accumulate the transcriptional regular RcsA, which is a positive regulator of capsular polysaccharide (exopolysaccharide) synthesis (*25, 26*). Increased exopolysaccharides has been shown to increase sensitivity to POL by increasing concentration of POL around the outer membrane in *Klebsiella* (*27*). We hypothesized that our TIG-resistant Lon mutants were overproducing exopolysaccharides, which in turn were making them more susceptible to POL. To test this, we randomly picked and grew three strains with the Lon mutation, three without the Lon mutation, and one with both the Lon and marR mutations on plates containing toluidine blue-O, congo red and ruthenium red. Toluidine blue-O binds to negatively charged polysaccharides, congo red binds to amyloid fibers like curli, and ruthenium red binds to acidic exopolysaccharides (*28*). The Lon mutants were preferentially stained by toluidine blue-O and congo red (Figure 2C), confirming that our Lon mutants overproduced exopolysaccharides. We did not see any difference between colonies grown on ruthenium red (data not shown). Based on these results, we propose that *lon*::IS186 mutants overproduce negatively charged exopolysaccharides which increase POL accumulation around the cells, rendering them hypersensitive towards the antibiotic (Figure 2D).

### MarR deactivation neutralizes Lon deactivation-driven polymyxin collateral sensitivity

If Lon mutations produced the POL CS phenotype in the TIG resistant cells, resistance to TIG that did not confer POL CS must have either occurred via a different pathway, or the CS effect of the Lon mutation may have been masked via secondary mutations. Candidate mutations that occurred frequently and preferentially in cells that remained neutral to POL were in the efflux regulators *acrR* and *marR* (Figure 2B) (Supplementary Figure 2), known to confer resistance to TIG (*29, 30*). This suggests that cells that bypass Lon mutations to achieve resistance via efflux upregulation avoid POL CS.

Fifty-five percent of the neutral strains also exhibited the Lon mutation (Figure 2B). How do strains that carry this mutation mask the CS phenotype? To answer this question, we narrowed our analysis to the SAGE mutants where we had an equal distribution of cells with CS and neutral phenotype sequenced (Supplementary Figure 2). While 100% of the neutral strains had a MarR mutation, 67% of them also carried the *lon*::IS186 mutation. Since deactivation of MarR allows upregulation of TIG and POL efflux (*22, 23, 31*), we hypothesized that deactivation of MarR on a *lon*::IS186 background neutralizes the CS phenotype associated with Lon mutants (Figure 2D). To test this, we constructed the pUC57-marR plasmid and introduced it into four POL-neutral strains that showed both the marR and *lon*::IS186 mutations. Strain 3 showed a 2 bp deletion mutation and strain 5 introduced a premature stop codon in marR (Table 1) which should both significantly reduce or completely abolish marR activity (*32*). Mutation in the 104th amino acid that changes the glycine carried by strain 4 (Table 1) has been associated with reduced marR activity (*33*). The AAGGCTGG duplication causes a frameshift mutation which should also affect marR activity (Table 1) (*34*). Introduction of pUC67-marR converted three of the four strains from neutral to CS to POL, and increased susceptibility in the wildtype strain (Table 1). This suggested that cells that mutated MarR gained the ability to resist POL at a magnitude large enough to neutralize CS due to the *lon*::IS186 mutation. Reintroduction of MarR on a plasmid then reduced efflux levels, reverting the effects of the MarR mutation.

**Table 1:** Changes in POL MIC after introduction of the pUC57-marR plasmid in strains with different MarR mutations.

| Strain | MarR allele | POL MIC (µg/mL) |
| --- | --- | --- |
| WT | - | 0.25 |
| TIG-SG-N-3 | Δ2 bp coding (428-429/435 nt) | 0.25 |
| TIG-SG-N-4 | G104S | 0.25 |
| TIG-SG-N-5 | Q117* | 0.25 |
| TIG-SG-N-7 | (AAGGCTGG)<sub>1→2</sub> | 0.25 |
| WT + pUC57 | - | 0.25 |
| TIG-SG-N-3 + pUC57 | Δ2 bp coding (428-429/435 nt) | 0.25 |
| TIG-SG-N-4 + pUC57 | G104S | 0.25 |
| TIG-SG-N-5 + pUC57 | Q117* | 0.25 |
| TIG-SG-N-7 + pUC57 | (AAGGCTGG)<sub>1→2</sub> | 0.125 |
| WT + pUC57-marR | - | 0.125 |
| TIG-SG-N-3 + pUC57-marR | Δ2 bp coding (428-429/435 nt) | 0.125 |
| TIG-SG-N-4 + pUC57-marR | G104S | 0.125 |
| TIG-SG-N-5 + pUC57-marR | Q117* | 0.125 |
| TIG-SG-N-7 + pUC57-marR | (AAGGCTGG)<sub>1→2</sub> | 0.125 |

MarR mutations cannot completely explain neutrality: a significant number of strains that carried marR mutations showed CS towards POL (Figure 2B) (Supplementary Figure 2). Additional mutations in these strains such as HldE (D-beta-D-heptose 7-phosphate kinase), GmhB (D-glycero-beta-D-manno-heptose-1,7-bisphosphate 7-phosphatase) and TamB (translocation and assembly module subunit) (Figure 2B) (Supplementary Figure 2) involved in LPS biosynthesis and maintenance (*22, 35, 36*) and may have played a role in POL sensitivity.

### Strong intra-class cross-resistance, but not collateral sensitivity, is prevalent in clinical E. coli

We analyzed antibiotic susceptibility data of 779 uropathogenic *E. coli* strains from the CANWARD surveillance study (*37*) to test for the presence of collateral effects (cross resistances and collateral sensitivities) of resistance in pathogenic *E. coli* (Supplementary Table 2). We identified strong cross-resistance between drugs of the same class, particularly between β-lactams (Figure 3A). Very little negative correlation (collateral sensitivity) was present in the dataset, which appears to be common in clinical datasets (*38*). The largest negative correlation of −0.05 was seen between omadacycline (OMC) and colistin (COL).

**Figure 3:**
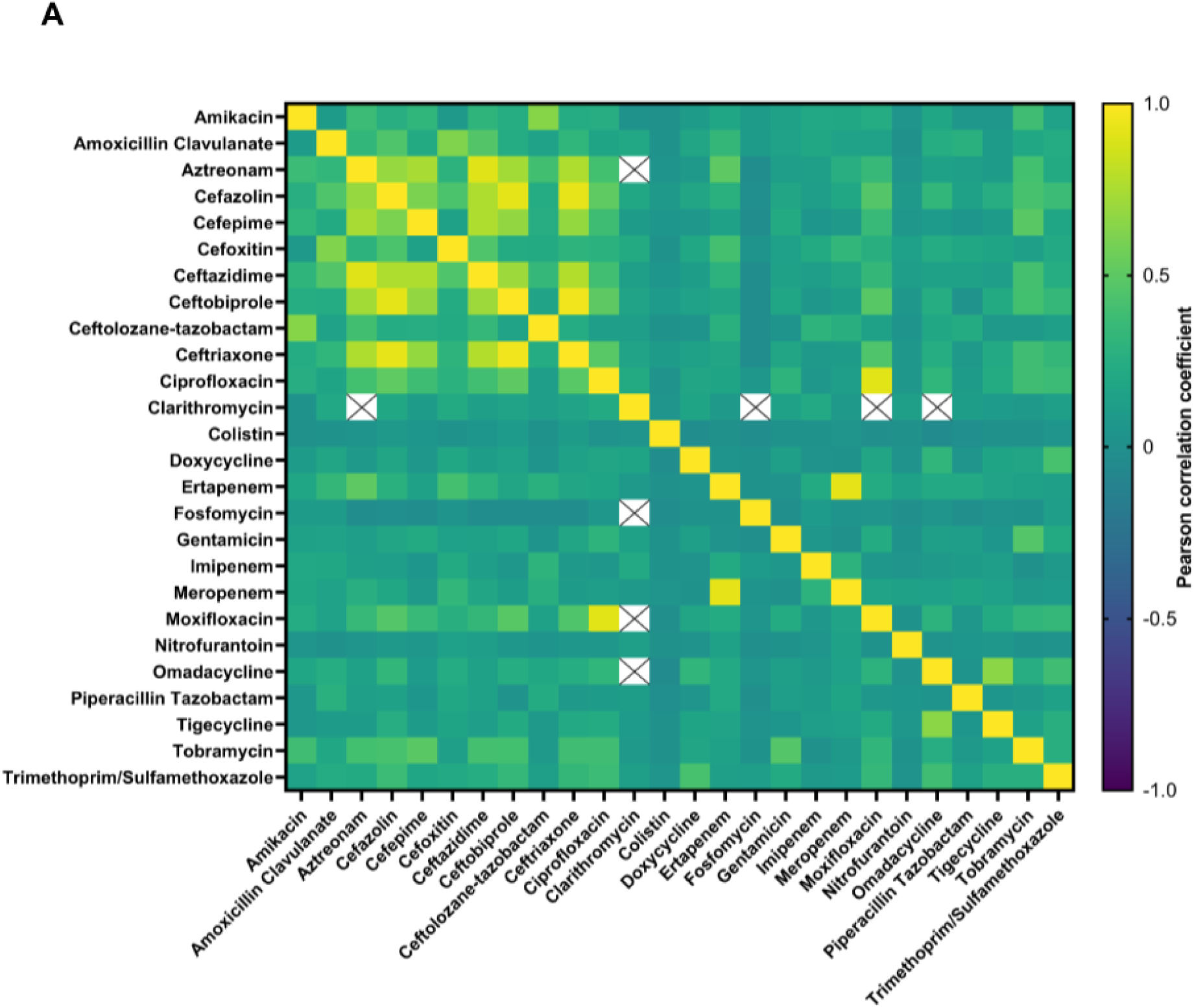
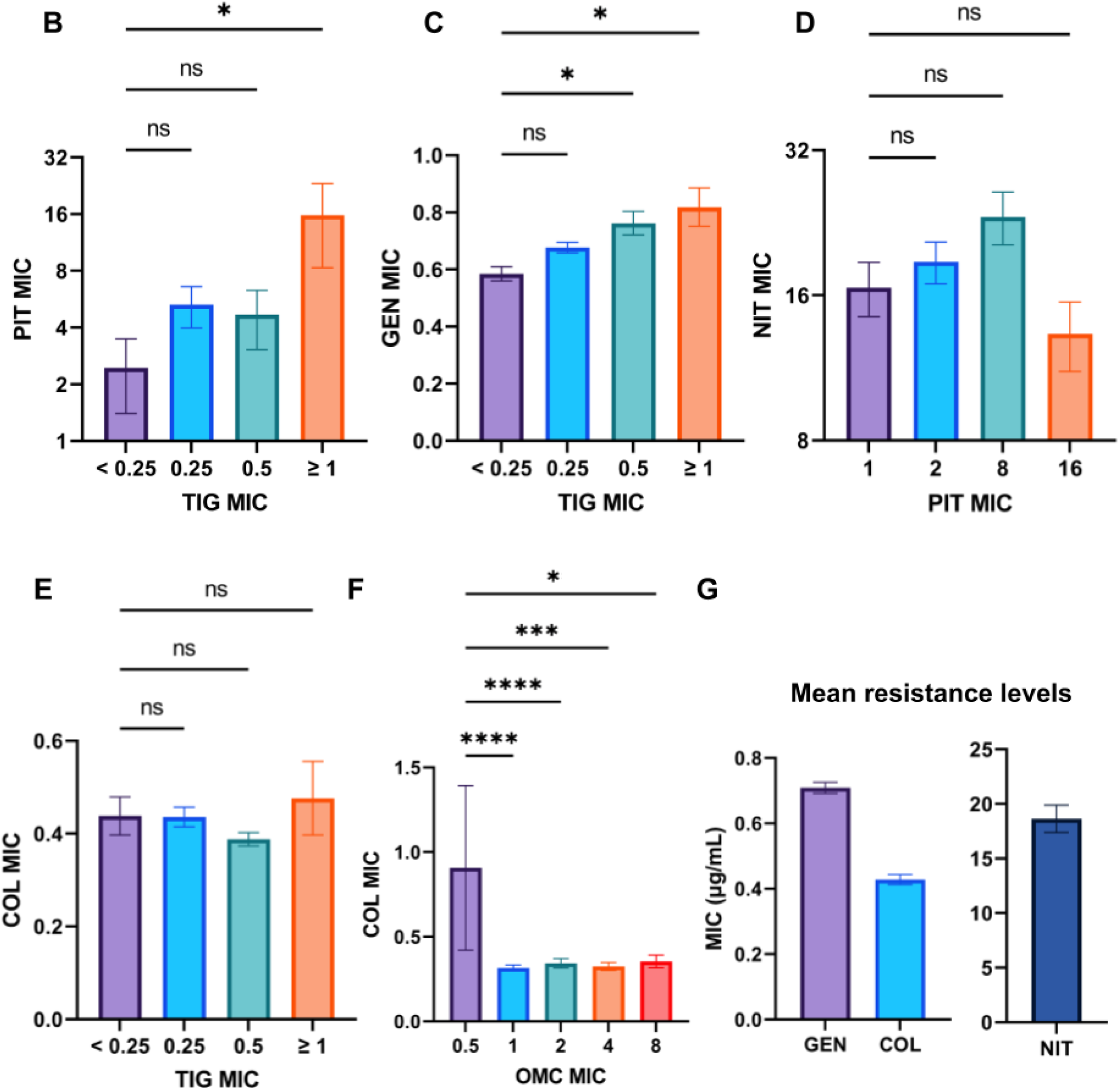
Uropathogenic *E. coli* antimicrobial susceptibilities reveals a rare CS relationship predicted by laboratory evolution. (A) Pearson correlation coefficients between MICs of each antibiotic with every other antibiotic in the dataset. A value of 1 denotes a perfect positive correlation (strong CR), 0 denotes no correlation, while a −1 denotes a perfect negative correlation (strong CS). (B) - (F) Relationship between resistances to drugs labelled on the x- and y-axes. *p<0.05, ***p<0.001, ****p<0.0001, one-way ANOVA with Bonferroni correction. (G) Mean resistance levels of antibiotics on the x-axis from the whole dataset.

### CS relationships rarely appear in clinical strains and their prevalence is best predicted by SAGE

We looked more closely at the clinical antibiotic susceptibility data to identify smaller changes in resistance. Specifically, we looked at the TIG-GEN, PIP-NIT and TIG-POL relationships to compare with our ALE results. PIP-STR was not included since STR was not among the drugs in the clinical dataset. Since our ALE adaptations only included chromosomal changes, it was important to limit our analysis to resistance conferred by chromosomal mutations instead of mobile elements. TIG is unaffected by Tet(M), tet(A), tet(B),tet(C), tet(D), and tet(K) mobile resistance determinants (*39*). Plasmid-encoded tet(X) genes that code for TIG inactivating enzymes and confer high TIG resistance with MICs ranging from 8-16 μg/mL are rare (*40, 41*), and our dataset did not contain TIG MICs >2 μg/mL. Chromosomal TIG resistance generally arises from increased efflux mediated by AcrAB-TolC in *E. coli* (*23*), and to further confirm that TIG resistance was largely chromosome mediated in our clinical dataset, we hypothesized that PIT (piperacillin/tazobactam) resistance should go up with increasing TIG resistance since PIP is susceptible to efflux. As expected, PIT resistance did increase with increasing TIG MIC in the clinical strains (Figure 3B).

Next, we looked at how GEN MICs varied with increasing TIG MIC. We removed GEN MICs > 4 μg/mL from the analysis because these MICs were more likely to be conferred by plasmid-borne aminoglycoside modifying enzymes (*42*). We observed a steady increase in GEN MICs with increasing TIG MIC among the remaining isolates (Figure 3C). From our ALE experiments, LQ and GP showed CS in over 25% of the strains, but SAGE in ∼4%, while CR was present in 17% of strains (Figure 1E) (Supplementary Figure 1).

NIT MIC changes with PIT resistance did not reach statistical significance, but showed an increasing trend up to 8 μg/mL of PIT resistance. From the ALE results, LQ and GP lineages showed CS in 17-25% of the strains with none from SAGE (Figure 1F) (Supplementary Figure 1). Over 40% of the SAGE lineages were cross-resistant to NIT (Figure 1F) (Supplementary Figure 1).

POL was not included in the clinical antimicrobial susceptibility data, so we examined the TIG-colistin (COL) relationship instead. Colistin and POL have very similar structures and near-identical activity *in vitro* (*43*). For the first time in the analysis, we see small but statistically insignificant reductions in COL resistance with up to 0.5 μg/mL of TIG MIC (Figure 3E).

During this analysis, we observed that mean COL resistance dropped significantly with resistance to another third generation tetracycline, omadacycline (OMC) (Figure 3F, G) (*44*). As OMC MIC increased from 0.5 to 1 μg/mL, COL MIC dropped ∼three-fold (Figure 3F). ∼86-96% of LQ and GP lineages and ∼39% of SAGE lineages showed CS to POL (Figure 1D) (Supplementary Figure 1).

## Discussion

In this study we investigated the repeatability of CS evolution in four reported drug pairs. With large sample sizes of 20-24 lineages we examined three different ALE platforms, testing the robustness of CS interactions to changes in evolutionary conditions and identifying possible ALE-specific biases. We found that SAGE allowed rapid evolution of high level resistance to antibiotics while still producing core resistance-conferring mutations comparable to other ALE platforms (Figure 1B, C) (Figure 2A) (Supplementary Figure 2). SAGE consistently produced lower frequencies of CS and higher cases of CR compared to the serial transfer and gradient plating-based methods (Figure 1D-G) in the TIG - GEN, PIP - NIT and PIP - STR drug pairs. This best matched antimicrobial susceptibility data from over 750 clinical MDR *E. coli* strains, in which CR and neutrality were abundant but significant CS relationships were almost entirely absent (Figure 3A). Importantly, we found no signs of CS between TIG - GEN, PIP - NIT and PIP - STR, and instead found evidence of cross-resistance (Figure 3C-E). This suggests that prediction of drug CS relationships from ALE experiments must be performed carefully. If CS is rare in clinical strains, ALE platforms with similar levels of CS will require a large number of replicates to model the clinical distribution. SAGE, with its low incidence of CS, appears well suited to this task.

Out of the four drug pairs tested, SAGE produced substantial CS in only one of them, the TIG - POL pair (∼39%; Figure 1D, H). From the clinical data, we observed that resistance to COL decreased with another third-generation tetracycline, omadacyline (Figure 3F) (*44*). The fact that this CS relationship appeared frequently in all three ALE platforms (Figure 1D) and held among MDR clinical strains suggests that this could be potentially exploitable to select against resistance, and that SAGE may be able to predict these important relationships.

We previously showed the presence of reciprocal CS between the TIG-POL drug pair (*20*), but the mechanism of CS was unknown. In this study we showed that tigecycline resistant Lon mutants produced increased extracellular polysaccharides, rendering them more susceptible to POL. From the ubiquity of the Lon mutations in our data and prior reports, it is likely that this mutation is the first step towards clinical TIG resistance (*23, 45*), and hence the COL sensitivity in clinical strains may be driven by the same mechanism. Cells may bypass the CS to POL by either increasing efflux through mutations in regulators like MarR, or by acquiring second step efflux regulator mutations after Lon deactivation (Figure 2B-D), suggesting the observed COL sensitivity may be transitory or that increased efflux may have undetected fitness costs.

Second step mutations in efflux regulators AcrR and MarR appeared more frequently in SAGE compared to LQ and GP (Supplementary Figure 2) and could partly be responsible for the low CS in SAGE lineages. In contrast to our findings, one study showed that accumulation of second step resistance mutations conferred collateral sensitivity over resistance to antibiotics (*46*).

However, in that study, efflux mutations were the first step mutations, with second step mutations conferring the CS they observed (*46*). This is in line with our and others’ findings that efflux mutations often confer broad-spectrum CR (*47, 48*). Another study showed how CS varies at a population level, with lineages that showed CS early during evolution getting replaced by mutants that acquired changes in genes with milder CS effects (*11*). Results from this study suggest that this can also occur at the strain level. Together, we suggest that the evolutionary timeline of CS cannot be generalized and must be studied on a case-by-case basis, possibly at the antibiotic and the bacterial species level.

The usefulness of CS has been debated, due in part to its dependence on the repeatability of evolution (*49*). Our results show that CS can indeed often appear at only low frequencies (Figure 1E-G), and is strongly linked to the specific mechanism by which cells become resistant. Our highly repeatable TIG-POL CS relationship was dependent on selection of the same *lon*::IS186 mutation across almost all sequenced lineages that exhibited CS. While this limits the number of CS interactions that may be worth pursuing, this also offers hope that robust CS that appear in complicated evolutionary landscapes in healthcare may be achieved via targeting collateral effects tied to readily accessible mutational hotspots.

From our attempts at evolving large sample sizes against multiple antibiotics via the different ALEs, we found that generating large numbers of resistant mutants was easier with SAGE (Supplementary Table 1), though the number of mutations per strain and the core resistance determinants selected for were similar to the other platforms tested (Figure 2A). This should facilitate ALE experimentation at scale in laboratories that do not have ready access to robotics.

Overall, we discovered a mechanism by which TIG resistance reliably conferred CS to POL by leveraging large scale laboratory evolution and showed that these biological effects were observed and reproducible in more than 750 clinical MDR *E. coli* strains. We highlighted the importance of large scale ALE experiments to generate robust profiles of collateral effects and showed that SAGE better predicts CS relationships that can help reduce antibiotic resistance in the clinic.

## Materials and Methods

### Bacterial strain and growth conditions

*E. coli* K-12 substr. BW25113 (WT) was grown aerobically at 37 °C with 250 rpm shaking in Muller Hinton (MH) broth. All evolved mutants were grown in MH broth supplemented with appropriate concentration of antibiotics.

### Susceptibility assays

MICs were performed using the EUCAST standard broth microdilution method (*50*).

### ALE experiments

SAGE, serial transfer and gradient plating-based evolutions were performed as described before (*18–20, 47, 51, 52*). For SAGE evolutions, the maximum concentration of antibiotics are listed in the table below. All SAGE plates were incubated for a fixed duration of seven days, which we found to be sufficient for cells to reach the end of the plates (*20, 53*). For serial transfers, evolutions were started at 1/8^th^ the WT MIC of the antibiotic by transferring 1 uL of overnight culture of WT bacteria into 99 uL of MH broth containing appropriate concentration of the antibiotic. Plates were incubated for 18-20 h, then 1 uL from all wells showing growth were transferred onto the 1/4^th^ MIC plate and so on, until the concentration listed in the table below was reached. Passaging beyond the listed concentration incurred significant loss of strains due to extinctions. For gradient plates, maximum antibiotic concentrations are listed below. Plates were made in square dishes (Falcon, Cat. No.: 351112) and were streaked towards the higher end of the antibiotic gradient after 18-20 h of incubation until growth was observed within the last grid of the plates.

| Antibiotic | Maximum concentration in SAGE |
| --- | --- |
| TIG | 5 µg/mL |
| PIP | 40 µg/mL |

| Antibiotic | Maximum concentration evolved to during serial transfers |
| --- | --- |
| TIG | 4x MIC, 1 µg/mL |
| PIP | 4x MIC, 4 µg/mL |

| Antibiotic | Maximum concentration in gradient plates | Days taken to reach end of plates |
| --- | --- | --- |
| TIG | 5 µg/mL | 5 days |
| PIP | 80 µg/mL | 5-7 days |

### Whole genome sequencing and analysis

Genomes were extracted using the Bio Basic genomic DNA kit (Cat. no.: BS624). Sequencing and variant calling was performed by Seqcenter (USA), on an Illumina NextSeq 2000. Variant calling was carried out using Breseq (80). NCBI reference sequence CP009273.1 was used for variant calling. Figure 2 and Supplementary Figure 2 were built using R and GraphPad Prism.

### Exopolysaccharide assay

MH + 1.5% agar was separately supplemented with 150 µg/ml Congo red, 40 µg/ml toluidine blue O or 40 µg/ml ruthenium red (AK Scientific) (*28*). All plates also contained 0.5 μg/mL of TIG. 10 uL of overnight cultures grown in MH broth + 0.5 μg/mL of TIG were spotted onto these plates.

### MarR complementation

The MarR fragment was synthesized based on its sequence available in the NCBI reference sequence CP009273.1, and inserted into the pUC57 MCS by Bio Basic Inc. The ligated plasmid was sequence verified before use in experiments. Cells were chemically transformed separately with the empty vector or pUC57-marR (*54*). We determined from separate experiments with this plasmid that IPTG induction was not required for sufficient marR production: cells transformed with this plasmid showed reduced efflux activity as measured by chloramphenicol resistance (*55*) but increasing concentrations of IPTG did not significantly change this resistance level (data not shown).

### Clinical strains and data analysis

Antibiotic susceptibility data of 779 uropathogenic *E. coli* strains was obtained from the CANWARD surveillance study (*37*). The Pearson correlation coefficients and MIC data extraction were performed using custom Python scripts.

## Acknowledgments

This study was funded by the Fonds de recherche du Québec – Santé (FRQS) (269182). FRC is supported by the Fonds de recherche du Québec – Santé (FRQS) (B2X). Veronica Banari was funded by DAAD Rise Worldwide during a Mitacs Globalink Research Internship. We thank L. Freeman for helpful discussions.

## Data and materials availability

WGS data is available under the NCBI sequence Read Archive BioProject: PRJNA1220725. All other relevant data is included in the main text and supplementary material. Strains are available upon request.

## Supplementary Figures

**Supplementary Figure 1:**
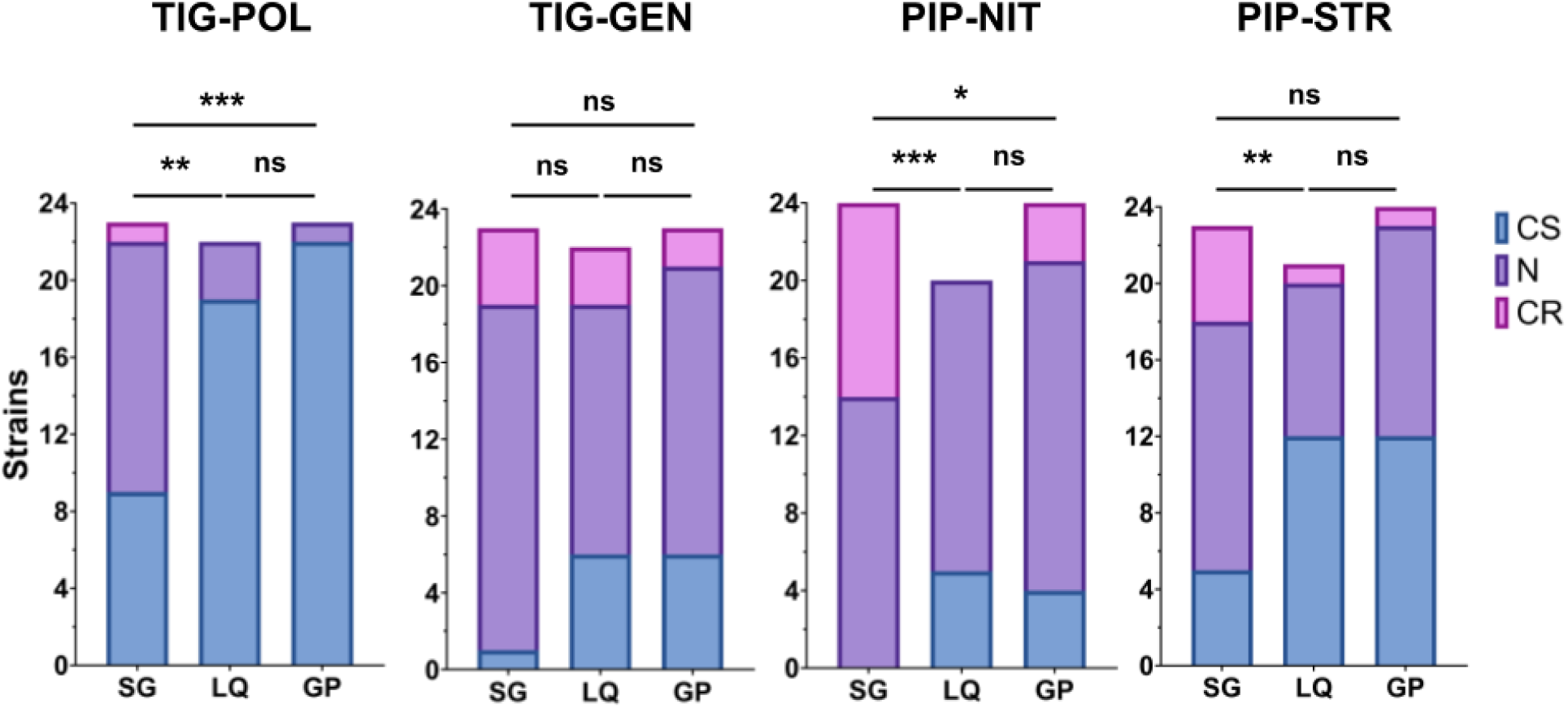
CS, N and CR distributions of the four antibiotic pairs tested, from each ALE platform. SG = SAGE. *p<0.05, ***p<0.001, ****p<0.0001, one-way ANOVA with Bonferroni correction. Statistical analyses were performed by comparing relative MICs of mutants from each platform.

**Supplementary Figure 2:**
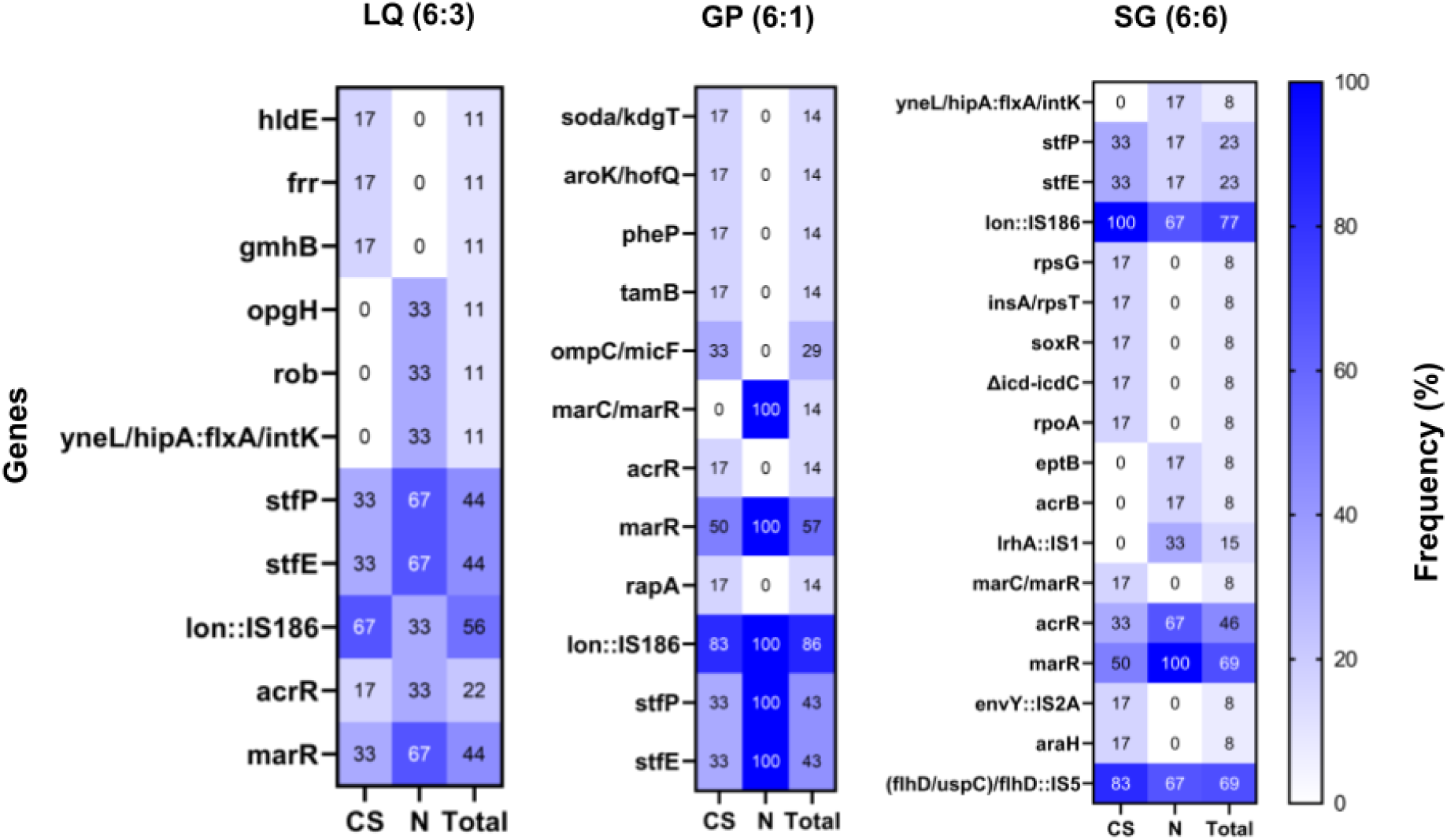
Frequency at which a mutation in the genes listed on the vertical axis appeared in each platform. Label on top denotes the platform while (6:3) = 6 CS strains and 3 N strains. For a detailed explanation of the x-axis and gene annotations, see the legend of Figure 2 (B). SG = SAGE.

---

**Supplementary Table 1:** Frequencies at which each platform generated resistant mutants against the different antibiotics used in the study.

| Antibiotics | ALE platform | Resistance frequency |
| --- | --- | --- |
| TIG | SAGE | 24 out of 24 evolutions |
|  | LQ | 22 out of 24 evolutions |
|  | GP | 23 out of 24 evolutions |
| PIP | SAGE | 24 out of 24 evolutions |
|  | LQ | 20 out of 24 evolutions |
|  | GP | 24 out of 24 evolutions |
| NIT | SAGE | 24 out of 24 evolutions |
|  | LQ | 3 out of 80 evolutions |
|  | GP | 24 out of 24 evolutions |
| CIP | SAGE | 23 out of 44 evolutions |
|  | LQ | 2 out of 24 evolutions |
|  | GP | Pilot plate showed no significant growth after 5 restreaks |

